# Rapid, modular, and cost-effective generation of donor DNA constructs for CRISPR-based gene knock-in

**DOI:** 10.1101/219618

**Authors:** Yi-Jiun Chen, Weikang Wang, Xiao-Jiun Tian, Daniel E. Lefever, David A. Taft, Jianhua Xing

## Abstract

CRISPR-based gene knock-in at endogenous sites is desirable in multiple fields such as quantitative studies of signal transduction pathways and gene regulation, synthetic biology, and disease modeling. Contrasting the knock-out procedure, a key step of CRISPR knock-in procedure relies on the homology-directed repairing (HDR) process that requires a donor construct as repair template. Therefore, it is desirable to generate a series of donor DNA constructs efficiently and cost-effectively. In this study, we developed a general Gibson assembly procedure that combines strengths of a Modular Overlap-Directed Assembly with Linkers (MODAL) strategy and a restriction enzyme based hierarchical framework. This procedure also allows fusing sgRNAs to the constructs for enhanced homology-directed repairing efficiency. Experimental tests on multiple constructs achieved from 3-8 folds of increase in assembly efficiency to high yield of constructs that failed to make with conventional Gibson assembly. The modularized procedure is simple, fast and cost-effective while making multiple constructs, and a computer package is provided for customized design.

## INTRODUCTION

In a cell, there are multiple intracellular signaling pathways to receive, transmit, and respond to intracellular and extracellular signals and regulate gene expressions. To investigate mechanisms of signal transduction, it is desirable to generate a series of knock-in mutants fused with reporters, such as fluorescence protein (FP), for tracking proteins dynamics [1, 2], and CRISPR knock-in is a method of choice. Unlike a knock-out procedure, a CRISPR knock-in process [3-5] requires additional donor DNA containing the specific knock-in sequence and repair templates, exploiting the mechanism of homology-directed repair (HDR) [6]. A typical donor DNA construct has four major components: the backbone vector, the insertion fragment (IF), such as FP, and the two homologous fragments (5’ and 3’ arms) on both sides of the insertion position of the target gene. A challenge is to generate a library of the constructs for multiple proteins in a targeted pathway efficiently and cost-effectively. Another challenge is to enhance the HDR efficiency for CRISPRCas9-induced precise gene editing. Some reported methods include using DNA nicks [7], suppression of KU70 and DNA ligase IV [8], and a HDR donor construct which is flanked by single guide RNA (sgRNA)-PAM sequences [9].

There are two categories of methods for assembling DNA fragments into a donor construct: restriction enzyme based assembly methods, such as BioBrickst [10], BglBricks [11] and Golden Gate [12], and sequence-independent overlap techniques, such as Circular Polymerase Extension Cloning (CPEC) [13], Sequence-Ligation Independent Cloning (SLIC) [14], Overlap Extension Polymerase Chain Reaction (OE-PCR) [15], and Gibson isothermal assembly [16, 17]. Gibson assembly is rapid and convenient because multiple fragments can be assembled in a defined order within a single-tube isothermal reaction. During an assembly reaction, an exonuclease creates single-stranded 3’ overhangs first that facilitate the fragments annealing through an overlap region at the end. Next, a polymerase fills in gaps within each annealed fragment and a DNA ligase seals nicks in the assembled DNA construct.

For increasing the efficiency of generating donor constructs, several simple and cost-effective cloning methods with high efficiency have been developed. Notably Casini *et al.* reported a Modular Overlap-Directed Assembly with Linkers (MODAL) strategy for synthetic biology [18]. This method uses in silico screening to design optimal linker sequences, which serves as overlapping sequences between DNA fragments for guiding assembly, and a modularized design allows repetitive usage of individual fragments. Guye *et al.* developed a hierarchical framework to generate complex gene circuits for genetic engineering in mammalian cells [19]. This approach combines usage of restriction enzyme digestion and Gibson assembly, and allows construction of synthetic gene circuits large in size.

In this work, we present a new procedure for generating a series of donor constructs efficiently through combining MODAL strategy, the hierarchical framework, and adding the sgRNA-PAM sequences for enhancing HDR efficiency. Our procedure can be applied to generate CRISPR knock-in mutants for tracing proteins in live cells.

## MATERIAL AND METHODS

### GC content calculations of all possible insertion sites in whole human genome

Assembled human reference genome (GRCh38/hg38) was downloaded from the NCBI database (GCF_000001405.34_GRCh38.p8_genomic.gbff), and split into individual GenBank files for every chromosome, and one corresponding to mitochondrial DNA. Analysis was performed on each chromosome individually. Alternate assemblies and unassembled reads were not included in the analysis. A combination of homebrewed scripts and Biopython [20] was used to parse the individual GenBank files and then extract the coding-domain-sequence (CDS) features for every file. For every CDS feature, the flanking genomic sequence (750 base pairs) up and downstream from both the start and stop codon were extracted. For each of these 4 regions the percentage of GC content was calculated by: (ΣG + ΣC) / ΣN × 100%, where N represents any type of the nucleobases. This percentage calculation was performed for every CDS feature in the assembled files that was subsequently turned into a list. Since there are potentially many different CDS entries for a given gene, which may or may not have different start/stop positions, the list was filtered such that only one CDS entry for a unique start/stop position was included. A command-line script is provided as a Supplementary Information named *chromo_gc_content.py* (requirement: Python 2.7 and python package Biopython (http://biopython.org/)).

### Computational screening of the linkers

We introduced a pair of linkers (a and b) into the IF vector (Fig. 1A). Unlike the original MODAL approach, we designed the linkers contain specific restriction enzyme sites, *Age*I at the 5’ end of linker a, and *Xho*I at the 3’ end of linker b, and six other restriction enzyme sites in the middle (See Supplementary Information). The linkers are needed for reducing the error rate of PCR amplification, increasing modularity of the procedure, and applying the hierarchical framework [19]. Therefore, we developed a computer programs to search the sequence space and select optimal linker sequences (Fig. 2). Briefly, first we generated all possible sequences of 30 base pair (bp) oligonucleotides, and subjected them to in silico screening to avoid adverse factors in PCR reaction, such as single-stranded DNA secondary structures, unbalanced GC content, potential of dimerization or hairpin structures, high salt adjusted melting temperature, GC clamp and nucleotide repeats, and high probability of cross dimerization. Next, we performed BLAST alignment screening to the survived linker sequences to select linkers with minimal sequence similarity to the whole human genomic library (Supplementary Fig. 1). Through the overall screening procedure, we identified 12 pairs of linkers (Table 1), and selected two pairs (pair 1 and 6) for subsequent experimental tests. See Supplementary Information for detailed screening information.

**Figure 1.**
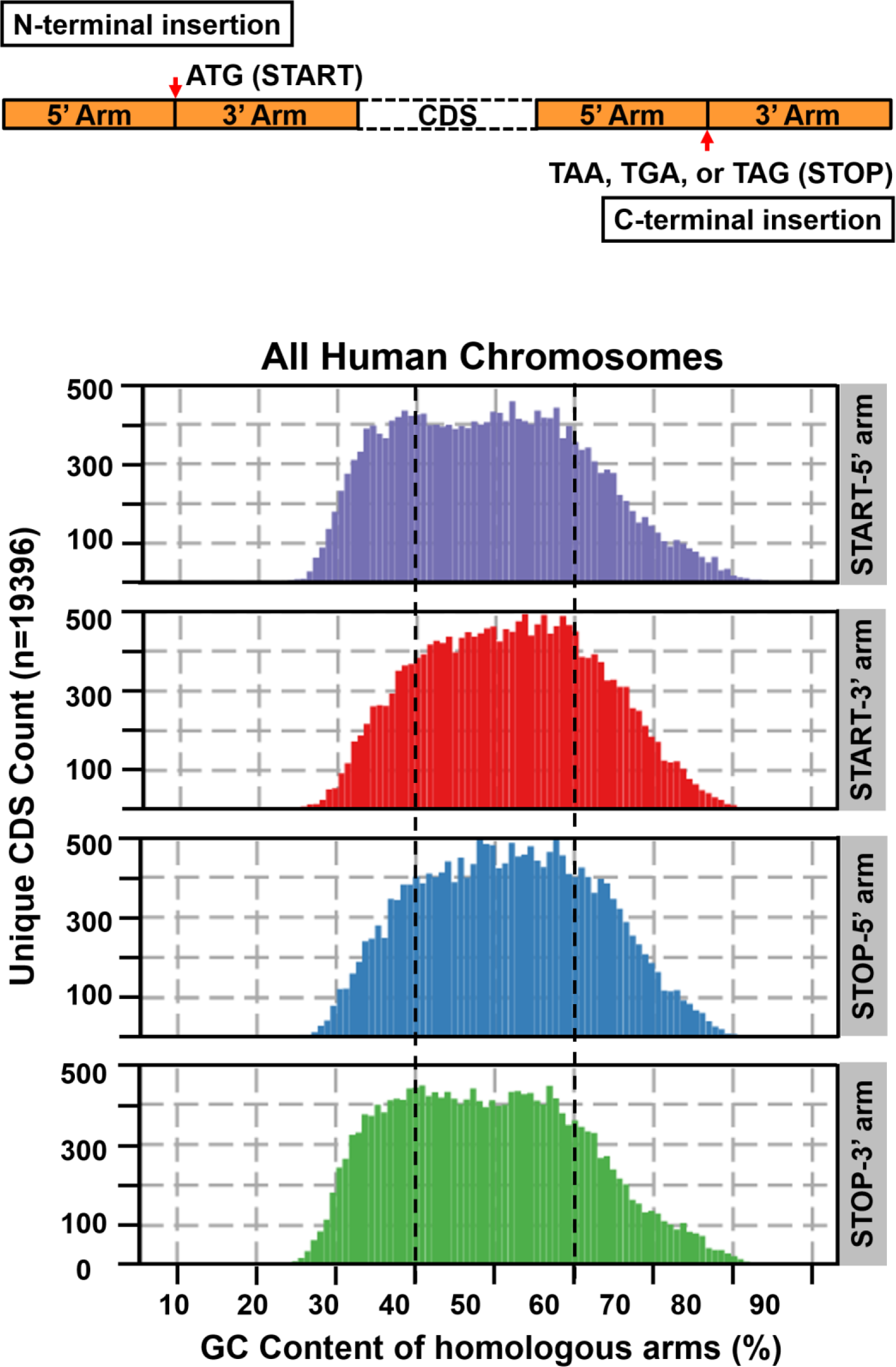

**Figure 2.**
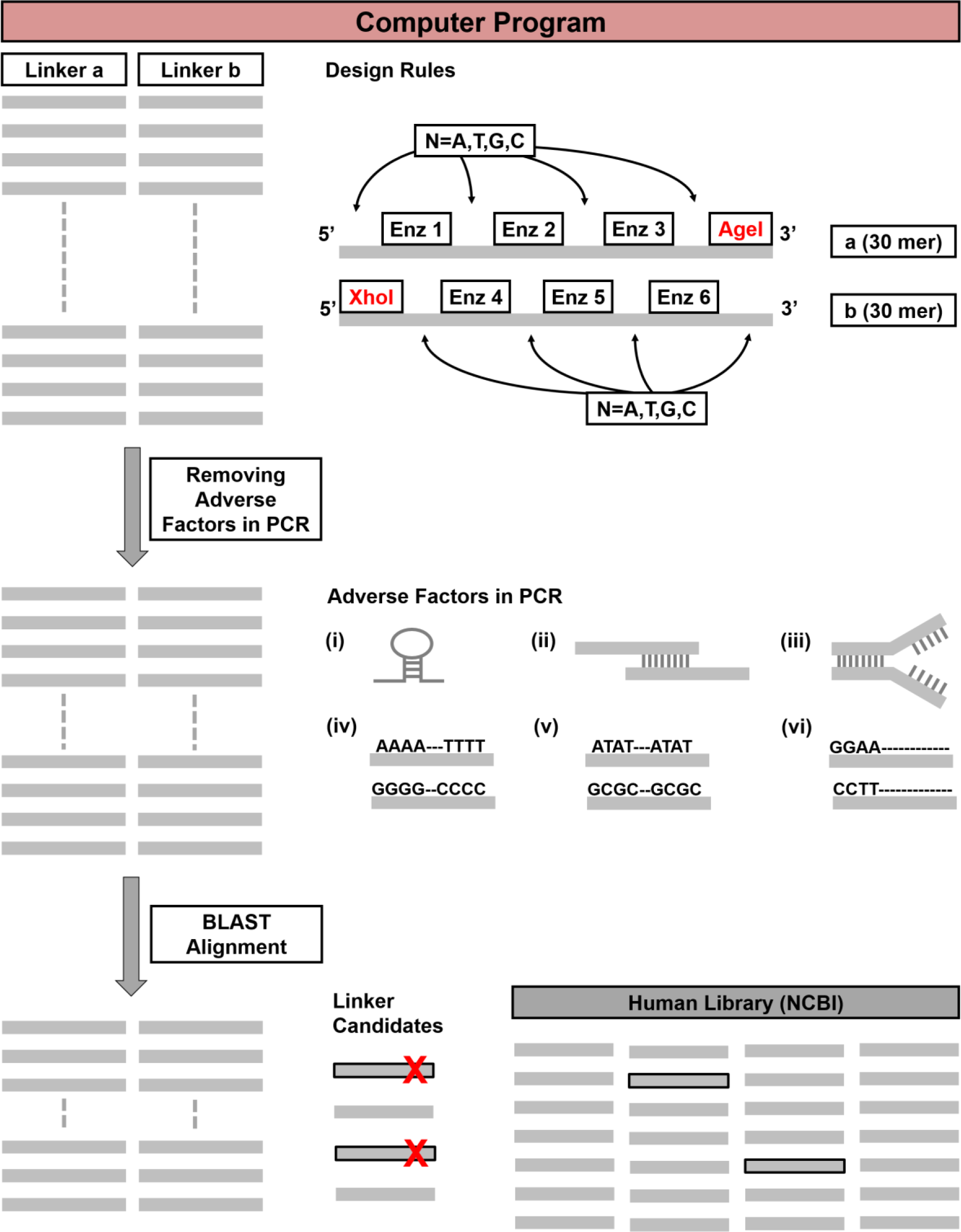

**Table 1.**
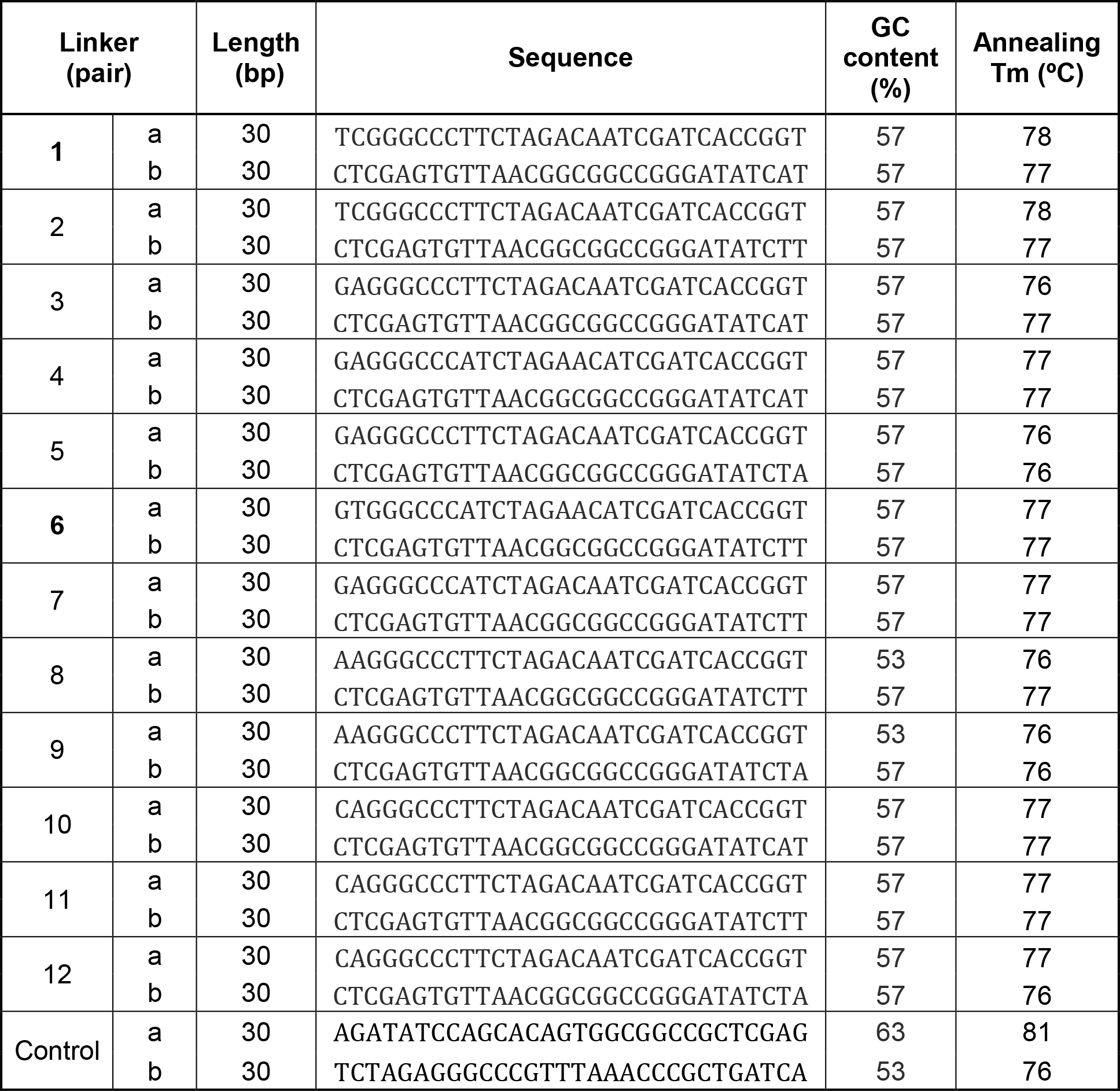
Top 12 ranked pairs of linkers given by in silico screening.

### Cloning and PCR methods

The donor constructs were generated by the Gibson assembly reaction (NEB#E2611). After transformation, we picked up colonies and grew them in LB medium with 50 ug/ml ampicillin. The GeneJET Plasmid Miniprep Kit (Thermo#K0503) was used for plasmid purifications. All of the primers used in this study are listed in Supplementary Table 2. For genomic DNA preparation, we used the QIAGEN DNeasy Blood & Tissue Kit (#69504) to extract the genomic DNA from T47D cells (ATCC#HTB-133). All homologous arms of knock-in targeted genes used in the assembly reactions were amplified by Q5 High-Fidelity 2X Master Mix (NEB#M0492) which is based on manufacturer's recommendations. All DNA fragments were purified by GeneJET PCR Purification Kit (Thermo#K0702) or GeneJET Gel Extraction Kit (Thermo#K0692). All restriction enzymes were purchased from New England Biolabs (NEB). For verification of donor constructs by PCR method, we used regular GoTaqG2 Flexi DNA polymerase (Promega#M7805) and the Deoxynucleotide Solution Set (dNTP, NEB#N0446). The FP vector was purchased from Addgene (pcDNA3-EGFP, #13031) and the linkers a and b were inserted to from IF vector which used in Fig. 2A. The mCherry construct was kindly provided by Dr. Rabin Lee. We amplified the mCherry with LoxP-Neomycin-LopX and inserted it into the pcDNA3 to form the IF vector which used in Fig. 3A.

**Figure 3.**
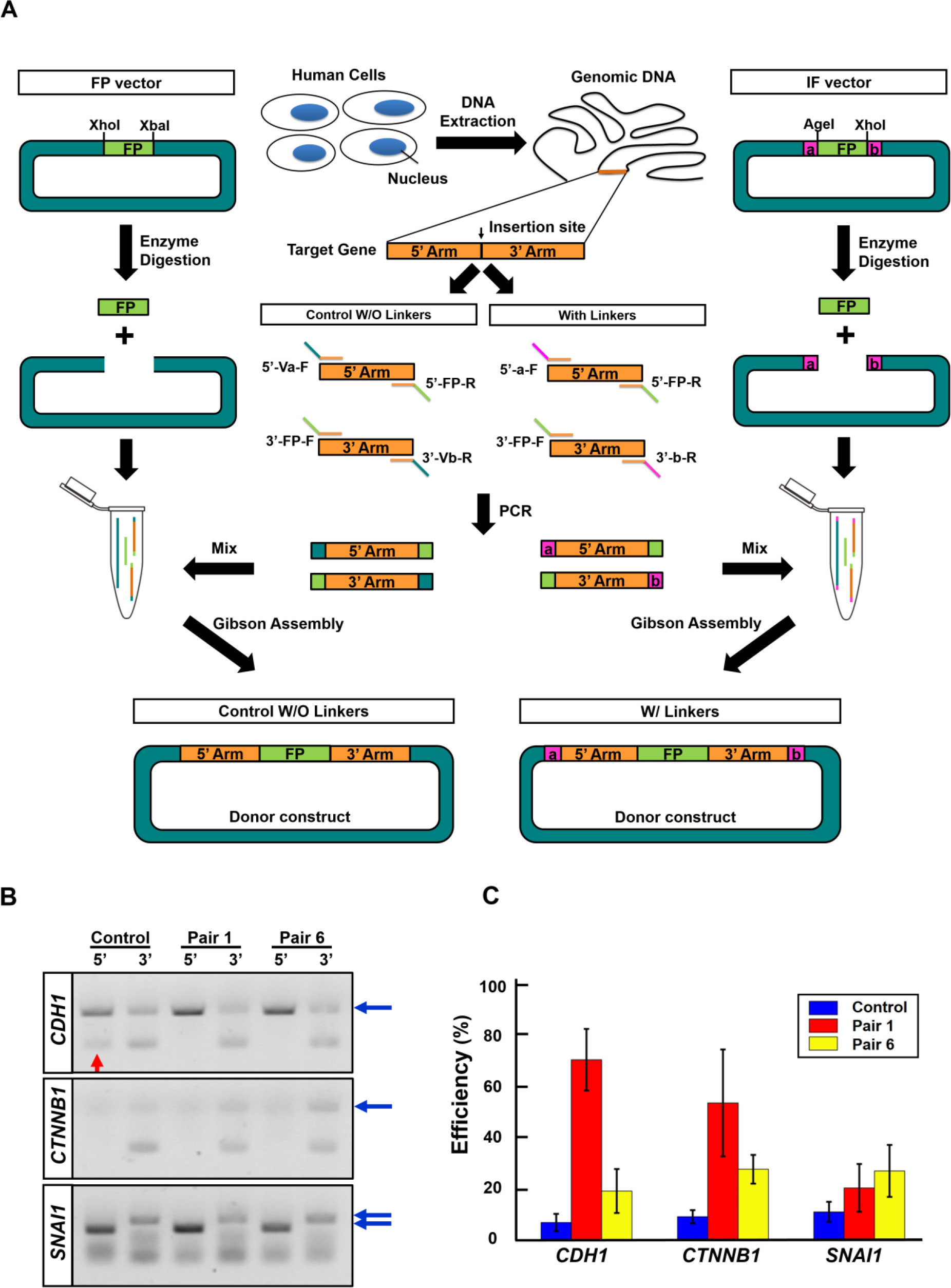

### Assembly efficiency measurements

All DNA fragments were purified first and mixed with NEBuilder HiFi DNA Assembly Master Mix (NEB#E2621) for one hour incubation time. The DNA assembly mixtures were transformed in competent E. coli DH5α and the transformation mixture were spread on the plate. Sixty colonies were picked for PCR and run DNA gel to check the insertion fragment. The efficiency was calculated as the number of colonies containing the desired construct divided by the total number of colonies.

## RESULTS

### High population of homologous arms with unbalanced GC content in human genes

A knock-in insertion site is commonly chosen as either the start codon at N-terminal or the stop codon at C-terminal of a target gene (Fig. 1, top). DNA synthesis is a direct and fast procedure for obtaining the 5’ and 3’ homologous arms. However, the DNA synthesis procedure is suitable for low-complexity fragments with few repeat sequences and homopolymeric region, and without unbalanced GC content [21]. Therefore, we first analyzed the whole human genome for potential insertion sites with flanking 750 base pairs (bp) region of start and stop codons. The result in Fig. 1 shows that for ∽45% of genes, one or both of the two flanking homologous fragments are GC unbalanced (>60% or <40%). This unbalanced GC content imposes challenge for getting the 5’ and 3’ arms through direct DNA synthesis.

### Twelve universal pairs of linkers were selected through in silico screening

Instead here we amplified the homologous arms of target genes from human genome directly through PCR. To enable the amplified homologous arms to be used in Gibson assembly, the primers need to contain two elements: the annealing sequences of the target gene for amplifying the homologous arm, and an overlapping region of 20–50 bp of shared sequence between fragments to direct Gibson assembly in a determined order [22]. In this study, we designed primers with a length of 60 bp, including 30 bp of annealing sequence for the target gene and 30 bp linkers for the overlapping region. To increase Gibson assembly efficiency and achieve modularity, we followed the MODAL strategy to design the linkers (Fig. 2). Different from the original MODAL approach, we selected optimal linker sequence against the human genome and required presence of multiple restriction enzyme sites (see Material and Method). We developed a computer program to generate sequences of 30 bp oligonucleotides, and subject all possible linkers to in silico screening to avoid adverse factors in PCR reaction. Next, the survival linkers needed to pass a BLAST alignment screening to minimize the identity between selected linkers and the whole human genomic library (Supplementary Fig. 1). Through the overall screening procedure, we identified 12 pairs of linkers (Table 1) and selected pairs 1 and 6 for subsequent experimental tests.

### The linkers increased Gibson assembly efficiency in generating multiple knock-in donor constructs

We selected genes related to the epithelial-to-mesenchymal transition (EMT) [23-26]. Inserting fluorescence protein (FP) sequences to the endogenous sites allows time-lapse live cell tracking of the spatial-temporal dynamics of EMT related proteins, such as silencing E-cadherin, and activating Vimentin and Snail1.

First, we generated a set of cloning vectors with the linkers and FP sequences (i.e. IF vector, Fig. 3). Then we chose three EMT-related genes *CDH1* (coding E-cadherin), *CTNNB1* (coding β-catenin), and *SNAI1* (coding Snail1) as examples. Due to high GC content and homopolymeric region, some homologous arms of these three genes cannot be synthesized via DNA synthesis. Therefore, we designed primers of these genes and amplified their homologous arms from human genomic DNA. The PCR results in Fig. 3B showed all 5’ and 3’ arms of three genes were successfully amplified with designed primers that fuse with overlapping sequence of the pair 1, pair 6, and control (without linkers), respectively. Specifically, the 5’ arm of *CDH1* using the control primer shows nonspecific band, which is absent while using pair 1 and 6 primers (red arrow). These results suggest that low sequence similarity between the linker sequences and the human genome may help on reducing non-specific amplification.

Next, we used EGFP as FP fragment for Gibson assembly efficiency test (Fig. 3C). We tested on the DNA construct for fusing *EGFP* to the C-terminal of gene *CDH1*. Compare to the control case, the linkers of pair 1 and 6 lead to ∽8 fold and ∽3 fold of increase of Gibson assembly efficiency, respectively. Similarly, we used the same linkers to test on the construct for fusing *EGFP* to the N-terminal of gene *CTNNB1*, whose product β-catenin is another major component of adherens junction. Compared to the control result, both linkers lead to ∽6 fold and ∽3 fold increase of Gibson assembly efficiency in generating CTNNB1 donor construct, respectively. For the construct fusing *EGFP* to the N-terminal of gene *SNAI1*, we achieved only ∽20% correct colony yield using the two pairs of linkers, which is still 2-3 folds increase of the efficiency compared to that of the control. These results indicate the in silico selected linkers indeed increased the Gibson assembly efficiency.

### High GC content of homologous fragments and removable SM severely reduce the Gibson assembly efficiency in generating the donor construct

Unlike the higher assembly efficiency in generating the donor constructs of *CDH1* and *CTNNB1*, the efficiency of that of *SNAI1* is much lower (Fig. 3C). It is unlikely due to unbalanced GC content of the linkers affecting the assembly efficiency [18], since the linkers that we used to amplify the three EMT-related genes have the same GC content (Table 1). Therefore we conjectured that unbalanced GC content of the homologous arms might affect the Gibson assembly. Indeed, the 5’ and 3’ arms of gene *SNAI1* have much higher GC content (72.8% and 60.5%), compare to those of the other two genes (Supplementary Table 3).

In addition, *SNAI1* has low expression in epithelial cells [27]. For such silent genes adding a removable selectable maker (LoxP-SM-LoxP) in the donor DNA is helpful for screening knock-in cells. We further tested the effect of adding a fragment LoxP-SM-LoxP with the FP to the N-terminal *SNAI1* donor construct (Supplementary Fig. 2A). The three LoxP sites (one on 5’arm and two on SM) interfere with the assembly process and lead to formation of incomplete DNA constructs (Supplementary Fig. 2B). This is in contrast with the low but practically acceptable efficiency we obtained for the corresponding construct without the SM (Fig. 3C). The LoxP-SM-LoxP sequence reduced the assembly efficiency to be undetectable, such as failed assembly, and imposed another severe challenge to Gibson assembly for the silent genes.

### A two-step procedure not only overcomes difficulties with unbalanced GC content and removable SM, but also allows adding sgRNAs for increased HDR efficiency

The results in Fig. 3C showed that the MODAL strategy alone cannot adequately resolve the issue of misassembly due to high GC content and the removable SM. To overcome these difficulties, we designed a two-step Gibson assembly procedure (Fig. 4A) to minimize the number of free fragment ends in each assembly step. The procedure took advantage of specific restriction enzyme sites (for *Age*I and *Xho*I etc) with low density in the whole human genome we included in the linkers (Supplementary Table 1), following the hierarchical framework [19]. These sites serve as restriction enzyme specific “unique cut” sites in the two-step procedure.

**Figure 4.**
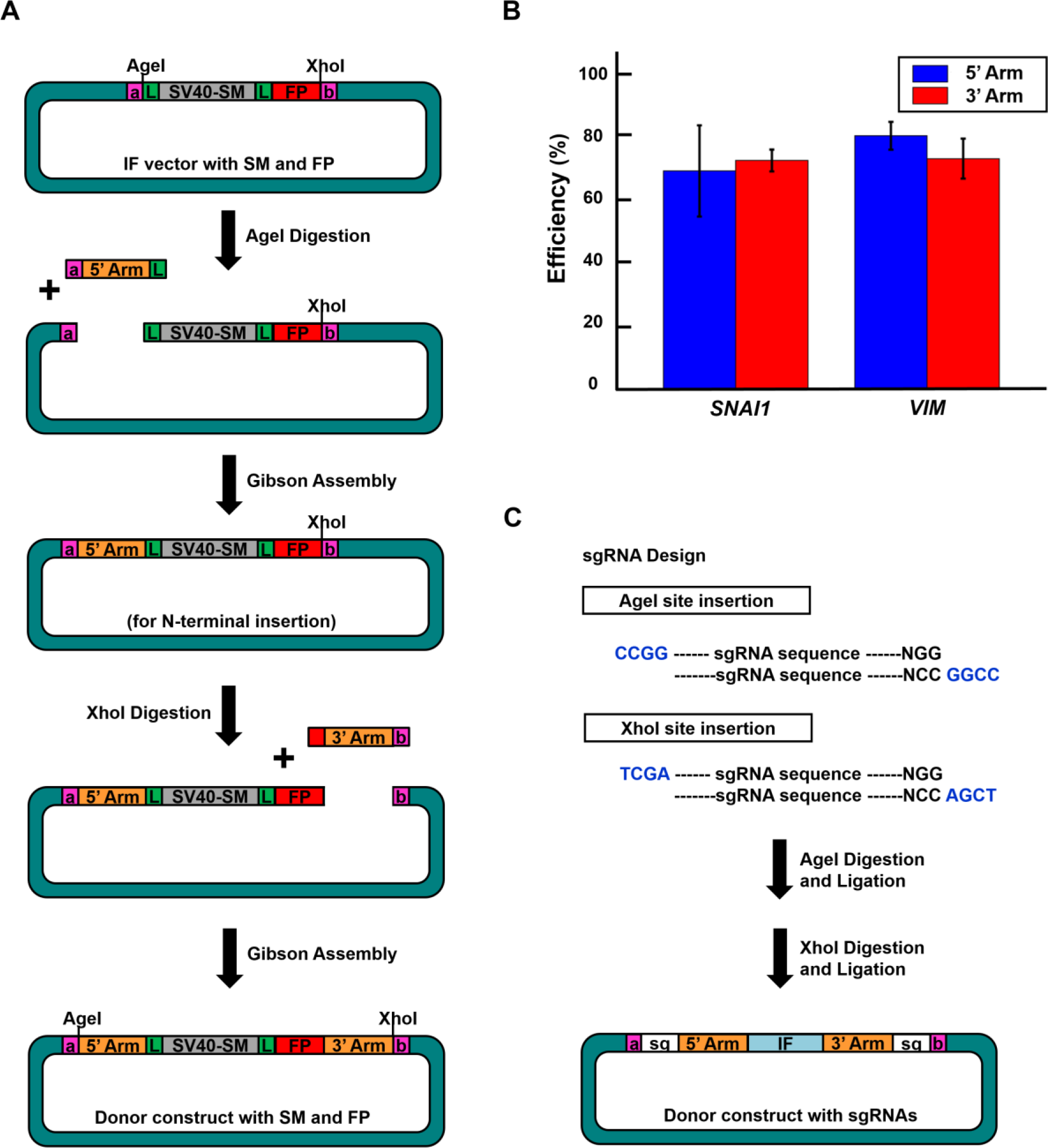

First we generated an IF vector that contains the linkers, LoxP-SM-LoxP, and FP fragments. After *Age*I digestion, we added the 5’arm and incorporated it to the vector through Gibson assembly. Next, we repeated the procedure but with *Xho*I digestion and 3’ arm incorporation. For the *SNAI1* donor construct with the SM and FP fragments, we achieved an assembly efficiency ∽70% for each of the 5’ and 3’ arms incorporation steps (Fig. 4B). We also achieved similar assembly efficiency on a donor construct for the *VIM* gene, a mesenchymal marker [23] that is a silent gene and has ∽70% GC content in the 5’ and 3’arms, respectively (Supplementary Table 3). These results indicated that the two-step procedure resolves the issues of low Gibson assembly efficiency due to high GC content and removable SM.

Furthermore, our linker design allows adding flanking sgRNA-PAM sequence at the donor construct, which has been shown to significantly enhance HDR efficiency [9]. Given that both of the *Age*I and *Xho*I sites are unique cutting sites in the donor construct, we designed sgRNAs that contain compatible ends of *Age*I or *Xho*I for the insertion (Fig. 4C). Following the similar two-step procedure of enzyme digestion, the donor construct with the flanking sgRNA-PAM sequences are inserted step by step through the enzyme digestion and ligation. We transfected the *CDH1-EGFP* construct with two sgRNA-PAM sequences into T47D cells and generated a knock-in cell line. The fluorescence images in Supplementary Fig. 3 show expected accumulation of EGFP at cell membrane.

## DISCUSSIONS

The CRISPR based gene editing techniques have opened multiple avenues in various fields of basic and applied science, but one impedance of wide applications of the technique in gene knock-in is that some generic properties of the required donor DNA constructs impose nontrivial technical challenges for DNA assembly. In this study, we developed an efficient procedure that combines strengths of three established approaches, the MODAL strategy, the hierarchical framework, and flanking sgRNA-PAM sequences. Experimental tests on selected DNA constructs showed that adding linkers lead to folds of increase in assembly efficiency, and a restriction enzyme-based two-step assembly procedure achieved high assembly efficiency for some difficult constructs with high GC contents and/or the LoxP-SM-LoxP sequence. Moreover, the two-step digestion and ligation further allowed insertion of flanking sgRNA-PAM sequences to donor construct for increased HDR efficiency. Both our one-step and two-step procedures are modularized to save the required time and cost.

While optimized for human genome in this work, the procedure and our accompanying software can be readily applied for other species. In addition, refinement of the selection criteria and expansion to whole sequence space search may further improve the performance of the linkers. The procedure can be further modularized by adding two flanking protein linkers N and C at the IFs (Fig. 5) following the designing rules reviewed by Chen et al [28]. Then the same homologous arms could be assembled with different FP (or RP) for the active genes or different SM and FP (or RP) for silent genes, i.e. repetitive usage of homologous arms. These multiple combinations provide more choices when generating donor constructs with different IFs for studying signaling pathways. We expect that this work and possible future refinement will facilitate CRISPR-based gene editing efforts in cell biology research, disease modeling, and synthetic biology studies in eukaryotic cells. For example, one can study functions of new genes by labeling and monitoring the temporal-spatial dynamics of the endogenous gene product [29]. One can generate models of fusion genes to study the physiological consequences in cancer development [30]. One can also insert regulatory elements at designated locations of a genome to manipulate the regulatory network for designed functions [31, 32]. Our developed procedure will greatly reduce the efforts needed on generating large libraries of DNA constructs in these studies.

**Figure 5.**
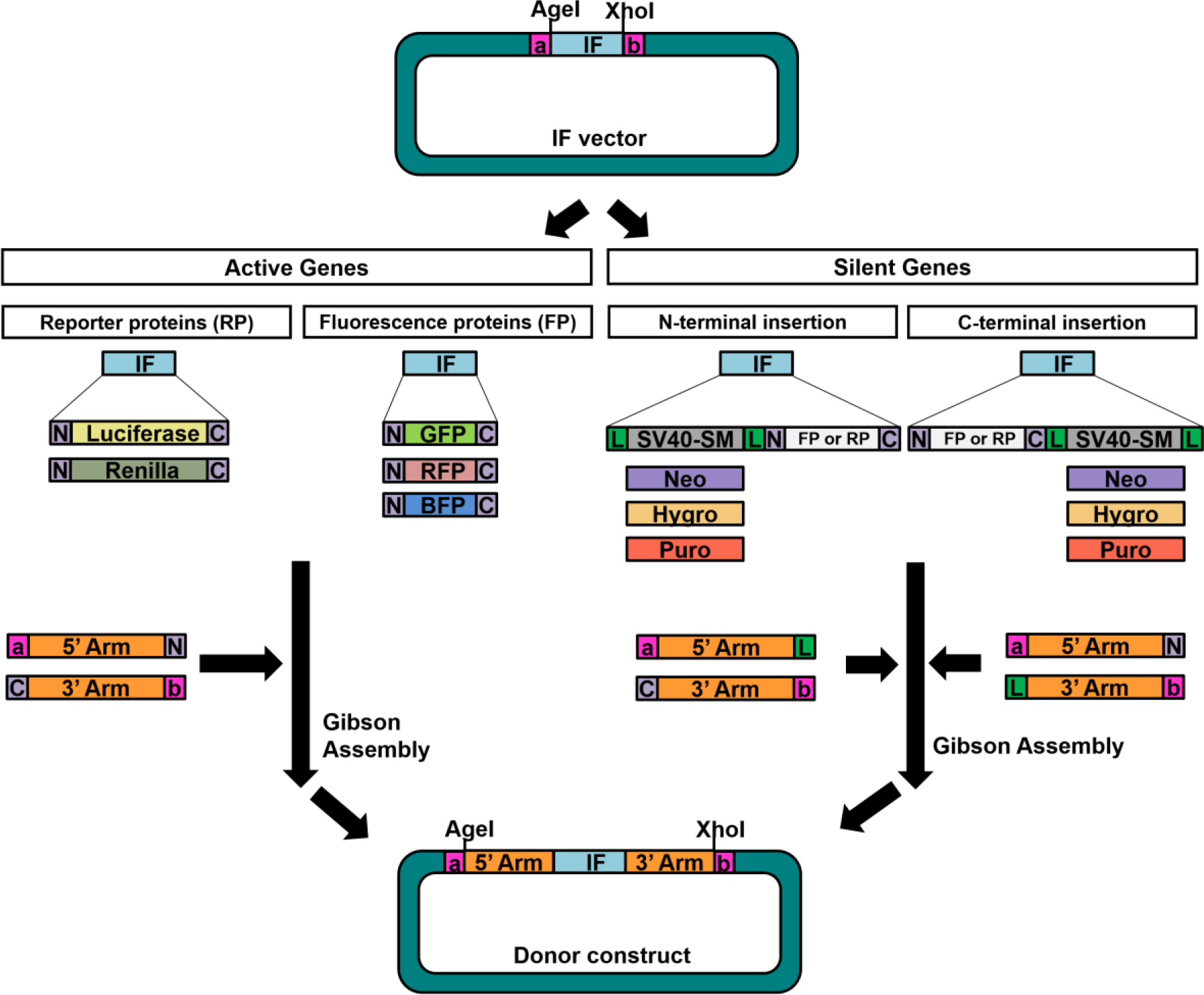

## ACKNOWLEDGEMENT

We thank Ms. Jingyu Zhang for helpful discussions.

